# Transferable representations of single-cell transcriptomic data

**DOI:** 10.1101/2021.04.13.439707

**Authors:** Ethan Weinberger, Su-In Lee

## Abstract

Advances in single-cell RNA-seq (scRNA-seq) technologies are enabling the construction of large-scale, human-annotated reference cell atlases, creating unprecedented opportunities to accelerate future research. However, effectively leveraging information from these atlases, such as clustering labels or cell type annotations, remains challenging due to substantial technical noise and sparsity in scRNA-seq measurements. To address this problem, we present HD-AE, a deep autoencoder designed to extract integrated low-dimensional representations of scRNA-seq measurements across datasets from different labs and experimental conditions (https://github.com/suinleelab/HD-AE). Unlike previous approaches, HD-AE’s representations successfully transfer to new query datasets without needing to retrain the model. Researchers without substantial computational resources or machine learning expertise can thus leverage the robust representations learned by pretrained HD-AE models to compare embeddings of their own data with previously generated sets of reference embeddings.

## Main

New developments in scRNA-seq technologies [1, 2, 3] are dramatically reducing the cost of experiments, facilitating the continual release of new scRNA-seq datasets and enabling the construction of large-scale, annotated reference atlases such as the Human Cell Atlas [4]. Despite this explosion in publicly available data, leveraging knowledge from previously studied scRNA-seq datasets to expedite the analysis of new datasets remains difficult since substantial nuisance factors of variation inherent in scRNA-seq measurements, such as dropout and transcriptional noise, can obscure biological signals of interest [5]. Moreover, combining measurements from multiple experiments is complicated by batch effects, i.e., systematic variations between datasets due to differences in experimental conditions or procedures. Batch effects are especially pronounced with scRNA-seq data, since different scRNA-seq protocols have unique sources of bias, sensitivity, and accuracy [6].

To address these challenges, several recent works [7, 8, 9, 10, 11, 12, 13] propose data integration methods that produce denoised low-dimensional representations (*embeddings*) of scRNA-seq data. However, these methods are not designed to integrate new query datasets with a previous reference set of embeddings without making users rerun the entire integration pipeline from scratch. This limitation is problematic; raw data from individual scRNA-seq experiments, even those within the same cell atlas, are often stored in different databases and in varying formats, necessitating a time-consuming data collection and preprocessing phase before integration can be performed (**Supplementary Figure 1**). Furthermore, current integration methods scale poorly in terms of computational cost and memory requirements [12] or require specialized hardware (e.g., GPUs), limiting their use to researchers with abundant computational resources.

As a response to these limitations, we introduce the Hilbert-Schmidt Deconfounded Autoencoder (HD-AE, **Figure 1a**), a deep learning approach based on the widely used autoencoder architecture [14] for learning denoised embeddings of scRNA-seq data. To ensure that samples’ embeddings are independent of their batches of origin, we train HD-AE using a loss function that penalizes the Hilbert-Schmidt Independence Criterion (HSIC) [15], a nonparametric measure of statistical independence, between samples’ embeddings and their batch labels (**Methods**). Removing batch information from the latent space would normally make it difficult for the autoencoder to reconstruct the data faithfully while also preserving true biological structure in the latent space; to mitigate this issue, when training HD-AE, we pass samples’ batch labels to the decoder so that batch-specific transformations can be learned to reconstruct the data accurately from the batch-effect-free latent space. *Unlike previously proposed scRNA-seq integration methods, pretrained HD-AE models’ representations successfully generalize to new batches of data not seen during training, even when those batches contain previously unseen cell types*. This lets researchers reuse previously trained HD-AE models off-the-shelf without needing to gather the original data or possess the computational expertise or hardware to train the models themselves from scratch (**Figure 1b**).

**Figure 1:**
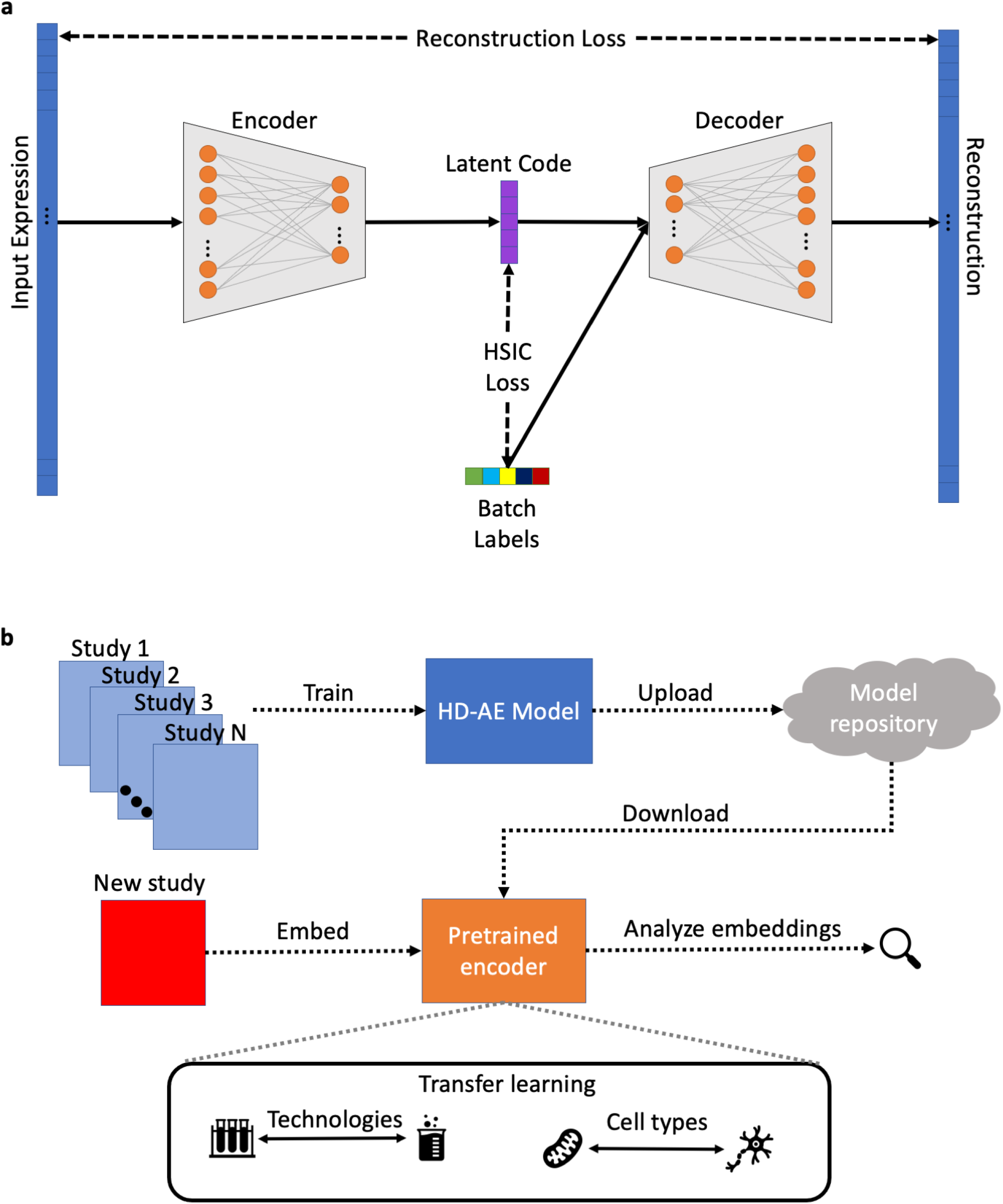
HD-AE models learn unified low-dimensional embeddings of scRNA-seq measurements originating from different experiments. **(a)** The HD-AE architecture. HD-AE encourages batch effect removal by penalizing the Hilbert-Schmidt Independence Criterion (HSIC) between samples’ latent representations and their batch labels (**Methods**). **(b)** A sample HD-AE transfer learning workflow. Researchers can download pretrained HD-AE models to embed their own data and compare with previously generated sets of reference embeddings.

We first applied HD-AE to construct a reference atlas of pancreas islet cell embeddings using three datasets, each sequenced using a different scRNA-seq protocol. Using UMAP [16] to visualize the raw data, we confirmed that it was clearly separable by batch, even for cells with the same cell type label (**Figure 2a**). After training an HD-AE model and embedding the data into the model’s latent space (**Figure 2b**), we observed that distinctions between batches were removed while cell types remained well-separated, indicating that embedding space variations were due to underlying biological differences rather than technical artifacts. To validate HD-AE’s transfer learning capabilities, we next used our pretrained model to embed a query batch of data collected using the CEL-Seq2 protocol, which was not used to generate any data seen by the model during training. We found that *embeddings of this query batch were well-integrated with training batch embeddings* (**Figure 2c**).

**Figure 2:**
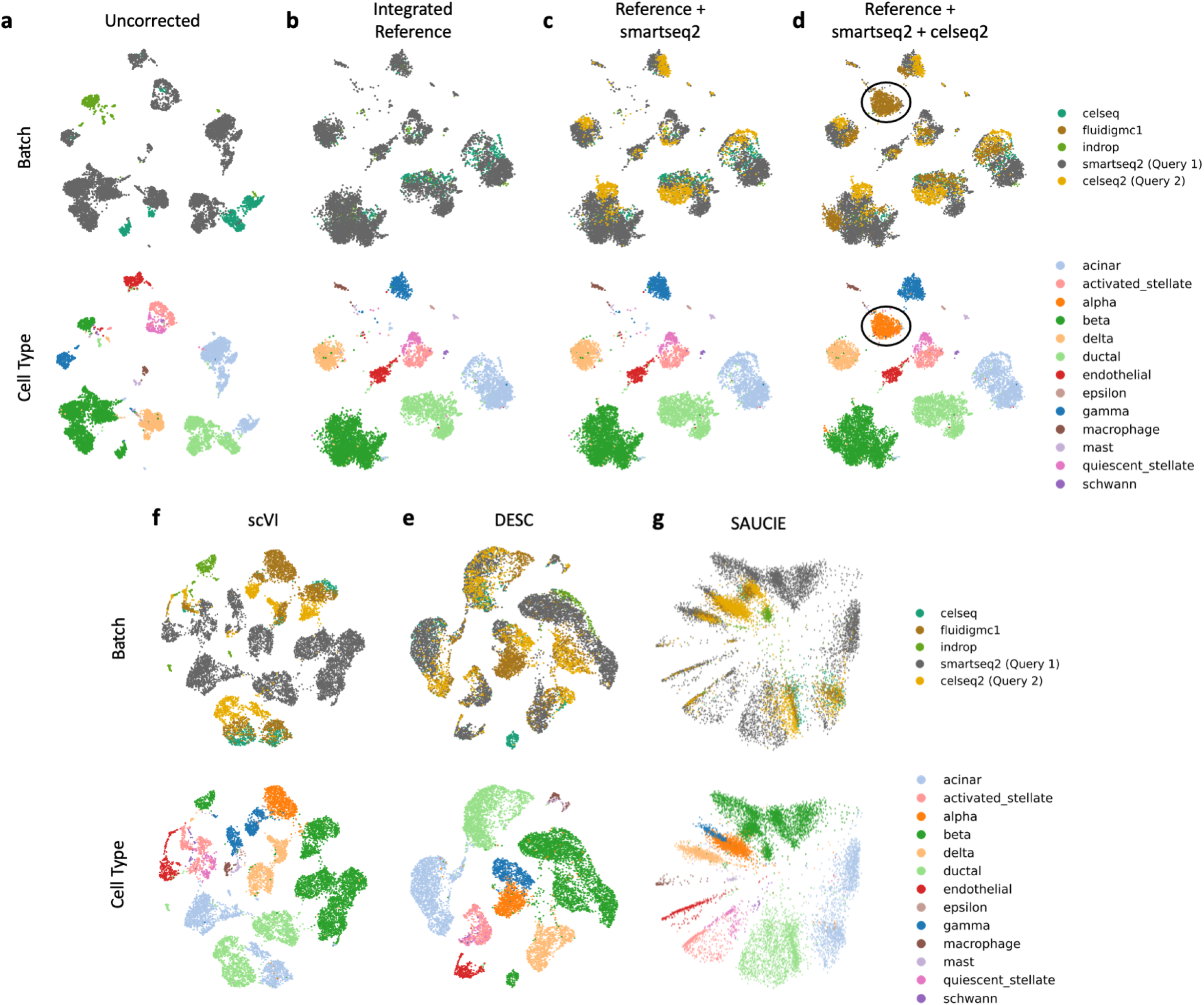
Unlike previously proposed deep learning embedding methods for scRNA-seq data, HD-AE’s representations generalize to new datasets at test time. **(a-b)** Three batches of pancreatic islet cells collected using different technologies from different labs before **(a)** and after **(b)** integration with HD-AE. **(c-d)** Querying the reference HD-AE model with two previously unseen batches of data. Black circles indicate cell types held out during training. **(e-g)** Embeddings produced by three previously proposed deep learning methods for scRNA-seq analysis using the same reference and query split as in **(a-d)**.

To further explore the robustness of our pretrained model, we also embedded a second query batch of data collected using the Smart-seq2 protocol (**Fig 2d**). To simulate a potentially more realistic scenario where new batches of data contain cell types not seen by the model during training, we included a cell type (alpha) in this second query batch that was held out during training. For cell types shared between the query and reference batches, we once again found that query batch cell embeddings were well-integrated with reference ones. Moreover, we found that the embeddings of the previously unseen alpha cells formed a distinct cluster well separated from other cell types. This behavior persisted for other choices of held-out cell types (**Supplementary Figure 2**). These results further indicate that *pretrained HD-AE models are able to filter out technical noise between different batches of data while preserving meaningful biological variations*.

We compared HD-AE’s transfer learning capabilities to those of three previously proposed deep learning methods for producing informative embeddings of scRNA-seq data: SAUCIE [17], scVI [11], and DESC [12]. None of these methods was originally designed for transfer learning. Nevertheless, their shared reliance on autoencoder-based architectures made it straightforward to adapt them for use in the transfer learning setting; we note here, however, that, limitations inherent in scVI forced us to disable its batch effect correction feature to use it for transfer learning (**Supplementary Note 1**). In these experiments, we used the same split between training and query batches as we did with HD-AE. Unsurprisingly, we found that scVI (**Figure 2f**) failed to produce well-integrated embeddings when not explicitly trained to correct for batch effects. Moreover, DESC (**Figure 2e**) and SAUCIE (**Figure 2g**) did not produce contiguous, well-separated clusters for individual cell types, possibly due to their choices of network architectures and loss functions (**Supplementary Note 2**). Only HD-AE produced clusters that were both well integrated between batches and well separated across cell types.

We next assessed HD-AE’s performance when applied to a dataset consisting of nine batches of peripheral blood mononuclear cells (PBMCs) collected using seven different technologies [18]. As we saw with the pancreas islet data, this dataset clearly separated by batch (**Figure 3a**). In this experiment, HD-AE was trained using seven of the batches; the remaining two (`10x Chromium v3` and `Drop-seq`) were held out as query batches. Once again, we found that batches were well-mixed and distinctions between cell types were preserved in the HD-AE embedding space (**Figure 3b**). Moreover, as we found with the pancreas data, our query batch and training batch embeddings were well-integrated (**Figure 3c-d**).

**Figure 3:**
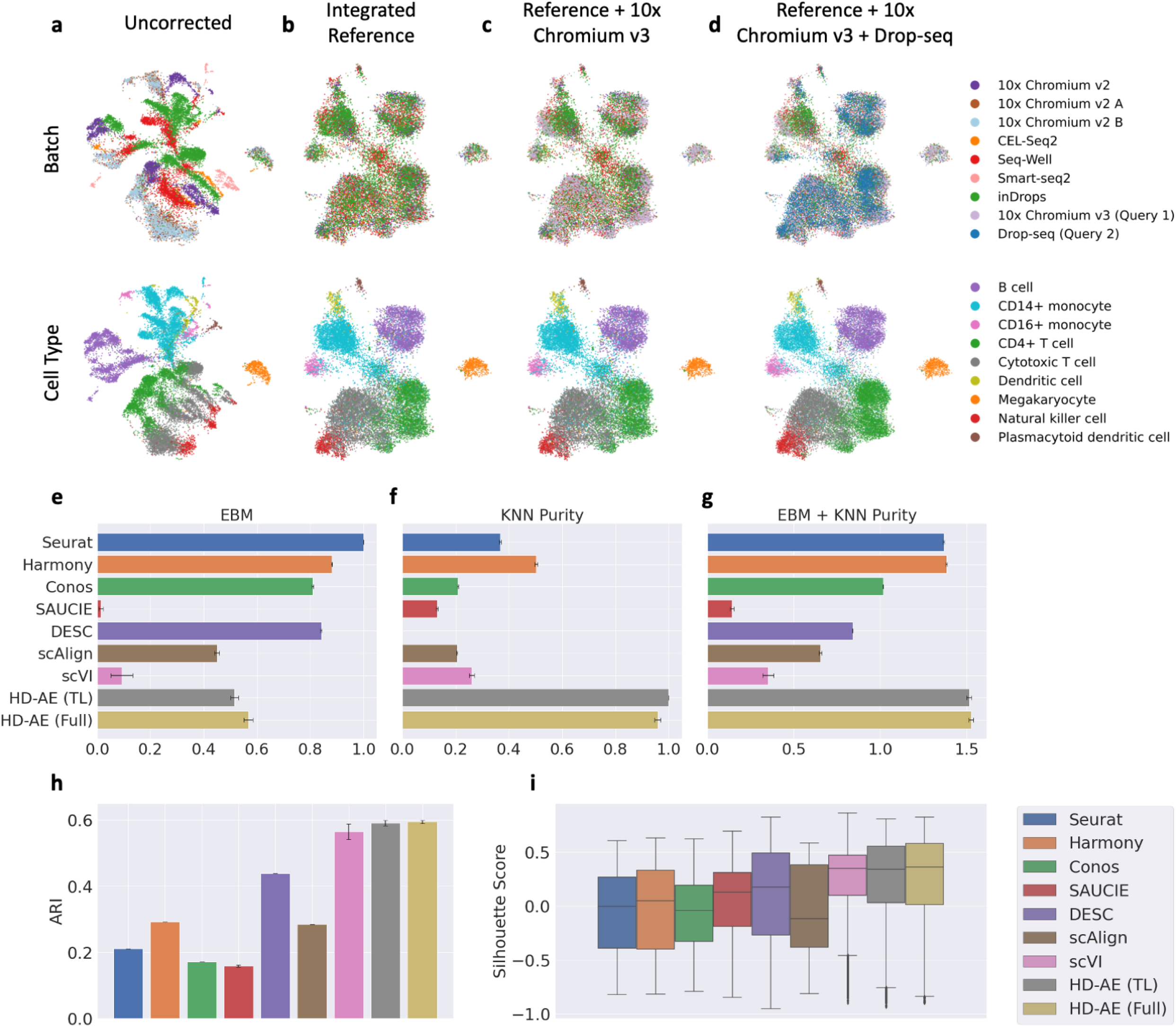
Transfer learning with HD-AE produces higher quality embeddings than previously proposed full integration workflow embeddings. **(a-b)** UMAP plots of seven batches of PBMC data before **(a)** and after **(b)** integration using HD-AE. **(c-d)** Querying the reference HD-AE model with two previously unseen batches of data. **(e-g)** Metrics for evaluating integration performance. Entropy of batch mixing (EBM, **(e)**) quantifies cell mixing across batches, while *k*-nearest neighbors purity (kNN purity, **(f)**) quantifies the preservation of within-batch local structure. We also report the sum of these metrics (right) to evaluate how well each method balances the two properties. HD-AE (TL) refers to HD-AE trained with seven of the nine batches and embedding the remaining batches via transfer learning, while HD-AE (Full) refers to HD-AE trained with all nine batches. kNN purity and EBM were normalized to lie in the range [0, 1] to enable comparison across the metrics. For each method we report the mean and standard error across five random subsamples of the data. **(h-i)** Separation between cell types in the embedding space as quantified by the adjusted Rand index (**h**) and silhouette scores (**i**).

Using this dataset, we also compared the quality of HD-AE embeddings to “full integration” method embeddings (i.e., those from methods designed to integrate all batches of data at once rather than for transfer learning). In particular, we benchmarked HD-AE against seven previously proposed data integration methods: Seurat v3 [8], Conos [10], Harmony [9], scAlign [13], scVI [11], SAUCIE [17], and DESC [12]. Each baseline method was given access to all nine batches of data during training rather than only a set of reference batches. As an additional full integration baseline, we trained an HD-AE model using all batches during training. We report implementation details and hyperparameter choices for all methods in **Supplementary Note 3**. Qualitatively, we found that many baseline methods struggled with this dataset, with only scVI and HD-AE producing well-separated clusters for each cell type (**Supplementary Figure 3**).

We also computed a suite of quantitative metrics to evaluate the quality of each method’s embeddings. To effectively integrate data, a method must balance two potentially opposing goals. First, batches should be well-mixed after integration; that is, the set of *k*-nearest neighbors around a given cell should be balanced across different batches. To quantify this mixing, we computed the *entropy of batch mixing (EBM)* [7] for each method’s embeddings (**Methods**). A high EBM can be achieved by randomly mixing the data and disregarding biologically meaningful variations. Thus, our evaluation also considered how well local neighborhoods in individual datasets were preserved in the integrated space. To quantify the preservation of this structure, we computed the *k-nearest neighbors purity (kNN purity)* [19] for each method (**Methods**). A high purity score could trivially be achieved without performing any mixing between batches. Thus, we considered performance on both metrics when evaluating a given method. To compare across metrics, individual metric values were normalized to lie in the range [0, 1].

We report our results for individual metrics for a neighborhood size *k* = 50 along with the sums of both metrics to indicate how well each method balances the two (**Figure 3e-g**). We found that HD-AE outperformed all baseline methods when considering both metrics. This result persisted for varying values of *k* (**Supplementary Figure 4**). Remarkably, we found that *HD-AE’s performance in the transfer learning setting was nearly identical to its performance when provided all the data during training*.

To examine how well each method preserved true biological variations, we also quantified how well different cell types clustered after integration. For each method we calculated the adjusted Rand index (ARI) to assess agreement between ground truth cell type annotations and cluster labels assigned by the Leiden community detection algorithm [20] (**Methods**). We found that HD-AE outperformed all baseline methods on this metric (**Figure 3h**). We also plotted the distributions of silhouette scores (**Methods**) for each method (**Figure 3i**). Here, we found that HD-AE, even in the transfer learning setting, was only narrowly bested by scVI in terms of median silhouette score. Moreover, we once again found that HD-AE’s performance on these metrics was nearly unchanged between the full integration and transfer learning settings. Taken together, these results further demonstrate *HD-AE’s ability to learn high-quality transferable representations of scRNA-seq data*.

Finally, we used this dataset to explore how the number of training batches affects HD- AE’s generalization performance. To do so, we trained HD-AE models with varying numbers of training batches and evaluated the quality of their embeddings of the full dataset as before. Qualitatively, we found that providing more batches during training initially produced more mixing between batches and more compact, well-separated clusters for each cell type (**Supplementary Figure 5**), though this effect appears to have diminishing returns as the number of batches increases. For our quantitative metrics (**Supplementary Figure 6**), we found that HD-AE’s silhouette score and ARI performance did not vary considerably for different numbers of training batches; however, we did initially see sharp increases in our combined EBM and kNN purity metric as the number of training batches increased. In particular, HD-AE began to outperform our full integration baseline models for combined EBM and kNN purity when provided with only four of the nine batches during training. These results suggest that *HD-AE achieves most of its generalization potential when it receives only a small number of batches during training, potentially enabling the use of pretrained HD-AE models even for tissues or organisms that are less well-studied and for which less data is publicly available*.

This work introduced the HD-AE framework for producing transferable representations of single cell transcriptomic data. In experiments on pancreas islet cell and PBMC scRNA-seq measurements, we found that HD-AE produced well-integrated reference sets of scRNA-seq embeddings and that pretrained HD-AE models successfully generalized to new batches at test time, even when those batches contained previously unseen cell types. This advancement may enable researchers to leverage pretrained deep learning models to obtain embeddings of their own data for use in arbitrary downstream tasks without needing to undertake the burdensome and skill-intensive process of training the models themselves. As part of future work, we envision training HD-AE models on a variety of tissues from various organisms and distributing them for the benefit of the wider scRNA-seq research community.

## Methods

### Autoencoder Model

HD-AE extends the standard autoencoder architecture. An autoencoder consists of two networks: (1) an encoder network *f*_*ϕ*_: *X* → *Z* parameterized by *ϕ*, which maps from an input space *X* ∈ ℝ^*M*^ to a latent space *Z* ∈ ℝ^*D*^, and (2) a decoder network *g*_*ψ*_: *Z* → *X* parameterized by *ψ*, which maps the latent space representation of a sample back to the original input space. The goal of the encoder network is to learn to map a given sample to the latent space *Z* so that the decoder network can simultaneously learn to faithfully reconstruct the original sample from its latent space representation. Moreover, we assume that *M >> D*; therefore, the latent space *Z* acts like an information bottleneck, capturing the strongest sources of variation in the original data in order to perform accurate reconstructions. In our implementations, both subnetworks consist of fully connected layers with rectified linear unit (ReLU) activations between them. We measure reconstruction loss using mean-squared error, so we train our two networks to solve the optimization problem

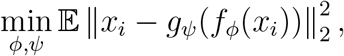

where the expectation is taken over our training data.

### The Hilbert Schmidt Independence Criterion (HSIC)

For random variables *X* and *Y* with probability distribution *p*_*XY*_, the HSIC measures the statistical dependence between the two. In particular, an HSIC of zero between *X* and *Y* is zero if and only if *X* and *Y* are independent, while a higher HSIC corresponds to a stronger level of dependence.

If 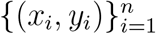 are independently and identically distributed samples drawn from *p*_*XY*_, the HSIC can be empirically estimated via

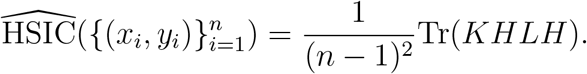

Here, *K*_*ij*_ = *k*(*x*_*i*_, *x*_*j*_) and *L*_*ij*_ = *l*(*y*_*i*_, *y*_*j*_) are Gram matrices for kernel functions *k* and *l*, respectively, where *k* and *l* must be universal kernels, a class that includes the widely used Gaussian and Laplacian kernels [15]. Moreover, *H* is a centering matrix, and Tr denotes the trace operator. See **Supplementary Note 4** for further details on the HSIC.

### HD-AE

HD-AE, an extension of the standard autoencoder model, was specifically designed to learn batch-effect-free latent representations. To do so, we added a regularization term to the autoencoder objective to minimize the empirical HSIC between latent representations and batch labels. Removing batch information from the latent space would usually complicate an autoencoder’s efforts to reconstruct the data faithfully while preserving true biological structure in the latent space; to mitigate this issue, when training HD-AE, we passed batch labels to the decoder so that batch-specific transformations could be learned to reconstruct the data accurately from the batch-effect-free latent space.

Suppose we have *n* total gene expression samples. Let *b*_*i*_ denote the batch label for the *i*th sample *x*_*i*_, let *B* denote the vector of batch labels for all samples, and let (by an abuse of notation) *f*_*ϕ*_(*X*) ∈ ℝ^*n×d*^ denote the matrix of latent representations of the dataset *X*. We then have the HD-AE objective function

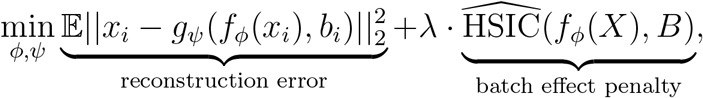

which we can optimize via stochastic gradient descent. In all our experiments Gaussian kernel functions with *σ*^2^ = 1 were used to compute 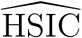.

### Datasets and Preprocessing

#### Pancreas Data

Our pancreas data came from the `panc8` dataset [21] provided in the SeuratData R package available at https://github.com/satijalab/seurat-data. We used data from the `celseq, fluidigmc1`, and `indrop` batches, as indicated by the `tech` field in the R object for training, and we used cells from the `smartseq2` and `celseq2` batches for testing generalization performance. We preprocessed the data by first filtering down the collection of datasets to the top 2000 highly variable genes as determined by the Seurat R package. For scVI, we used this filtered data directly; for other models, we normalized the data using the Seurat normalization workflow. For DESC, we also scaled the data after normalization using the `scale_bygroup` function from the DESC Python package.

#### PBMC Data

Our PBMC data came from the `pbmcsca` dataset [18] available in the SeuratData R package. For our transfer learning HD-AE model, we used data from the `CEL-Seq2, 10x Chromium (v2), 10x Chromium (v2) A, 10x Chromium (v2) B, Drop-seq, Seq-Well`, and `inDrops` batches, as indicated by the `Method` field in the R object for training. During preprocessing, we removed any cells with a cell type label of `Unassigned`; otherwise, our preprocessing workflow was the same as for the pancreas data.

### Evaluation Metrics

#### Entropy of Batch Mixing

Letting *c* denote the number of batches, the entropy of batch mixing (EBM) is defined as

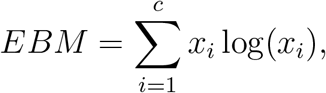

where *c* is the number of batches, *x*_*i*_ denotes the proportion of cells originating from a batch *i* in a given region, and 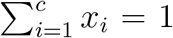. To assess the EBM for a given method, we followed a standard [11] estimation procedure: we randomly chose 100 cells, calculated “regional” EBM values for each cell using the batch proportions from the cell’s 50 nearest neighbors in the integrated space, and then averaged over the 100 regional EBMs.

#### kNN Purity

For a given batch, two similarity matrices were constructed. The first was computed using that batch’s cells’ gene expression values pre-integration; the second was computed using that batch’s cells’ representations in the integrated space. We then computed the ratio of the intersection of these matrices’ corresponding *k* nearest neighbors graphs over their union. We repeated this procedure for each batch in a dataset and reported the average of this statistic.

#### Silhouette Score

For a given cell *i*, the sillhouete score *s*(*i*) is defined as follows. Let *a*(*i*) be the average distance between *i* and the other cells in *i*’s cluster, and let *b*(*i*) be the smallest average distance between *i* and all other cells in a different cluster. The silhouette score *s*(*i*) is then

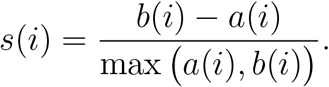

A silhouette score close to one indicates that *i* is tightly clustered with cells with the same ground truth label. A score close to -1 indicates that a cell has been grouped with cells with a different label.

#### Adjusted Rand Index

The adjusted Rand index (ARI) measures agreement between reference clustering labels and labels assigned by a clustering algorithm. Given a set of *n* cells and two sets of clustering labels describing those cells, the overlap between clustering labels can be described using a contingency table, where each entry indicates the number of cells in common between the two sets of labels. Mathematically, the ARI is calculated as

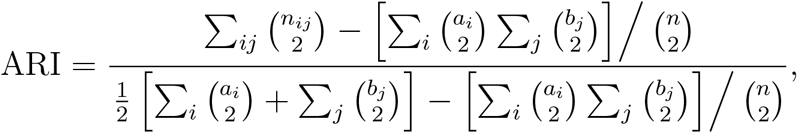

where *n*_*ij*_ is the number of cells assigned to cluster *i* based on the reference labels and cluster *j* based on a clustering algorithm, *a*_*i*_ is the number of cells assigned to cluster *i* in the reference set, and *b*_*j*_ is the number of cells assigned to cluster *j* by the clustering algorithm. In our experiments, we assigned cells to clusters using the Leiden community detection algorithm. Because the results of this algorithm depend heavily on its resolution hyperparameter, for each method we tried a number of resolution values in the range [0.5, 1.0] and reported the best resulting ARI for each method.

