## Supplementary Information for "Transferable representations of single-cell transcriptomic data"

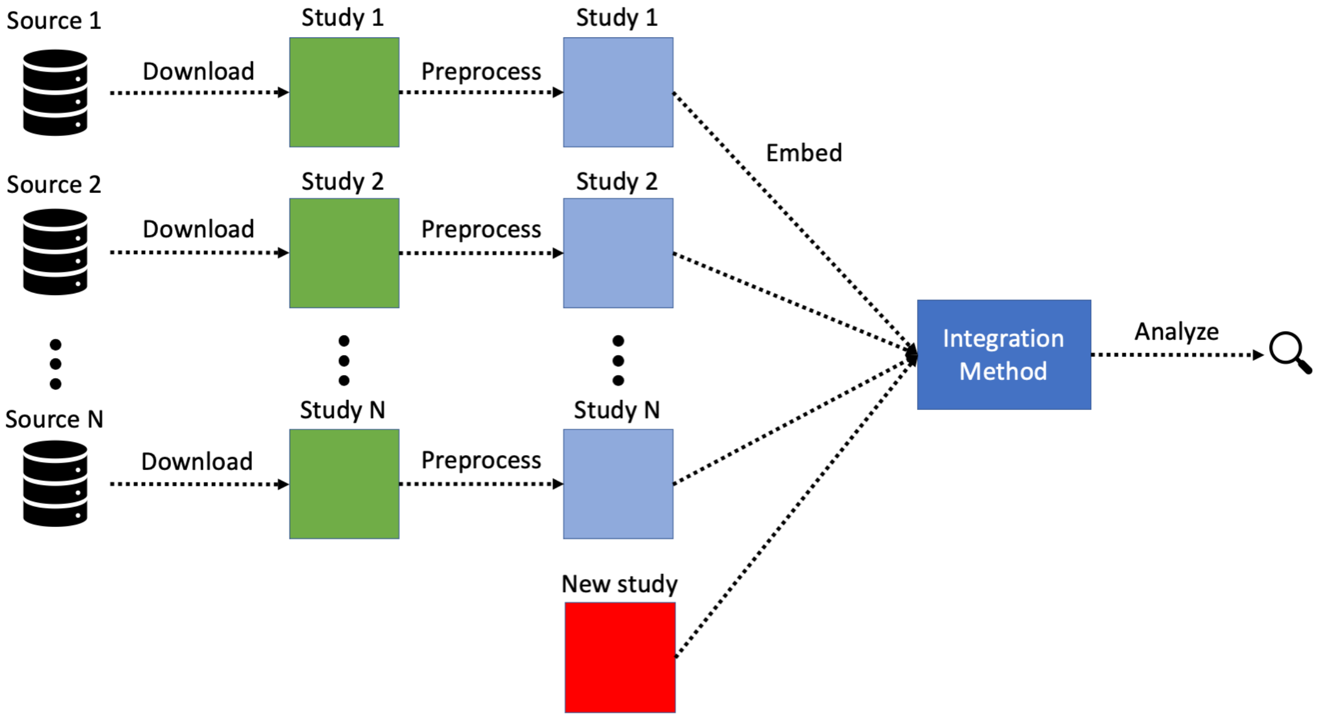


Supplementary Figure 1: Current workflow for integrating new datasets with previously annotated cell atlases. To use annotated atlas data to assist in the analysis of a new dataset, a researcher must: (1) retrieve each individual dataset making up the atlas, likely involving searching through multiple data stores, and (2) preprocess each dataset individually, which likely requires working across multiple file formats and/or data structures.

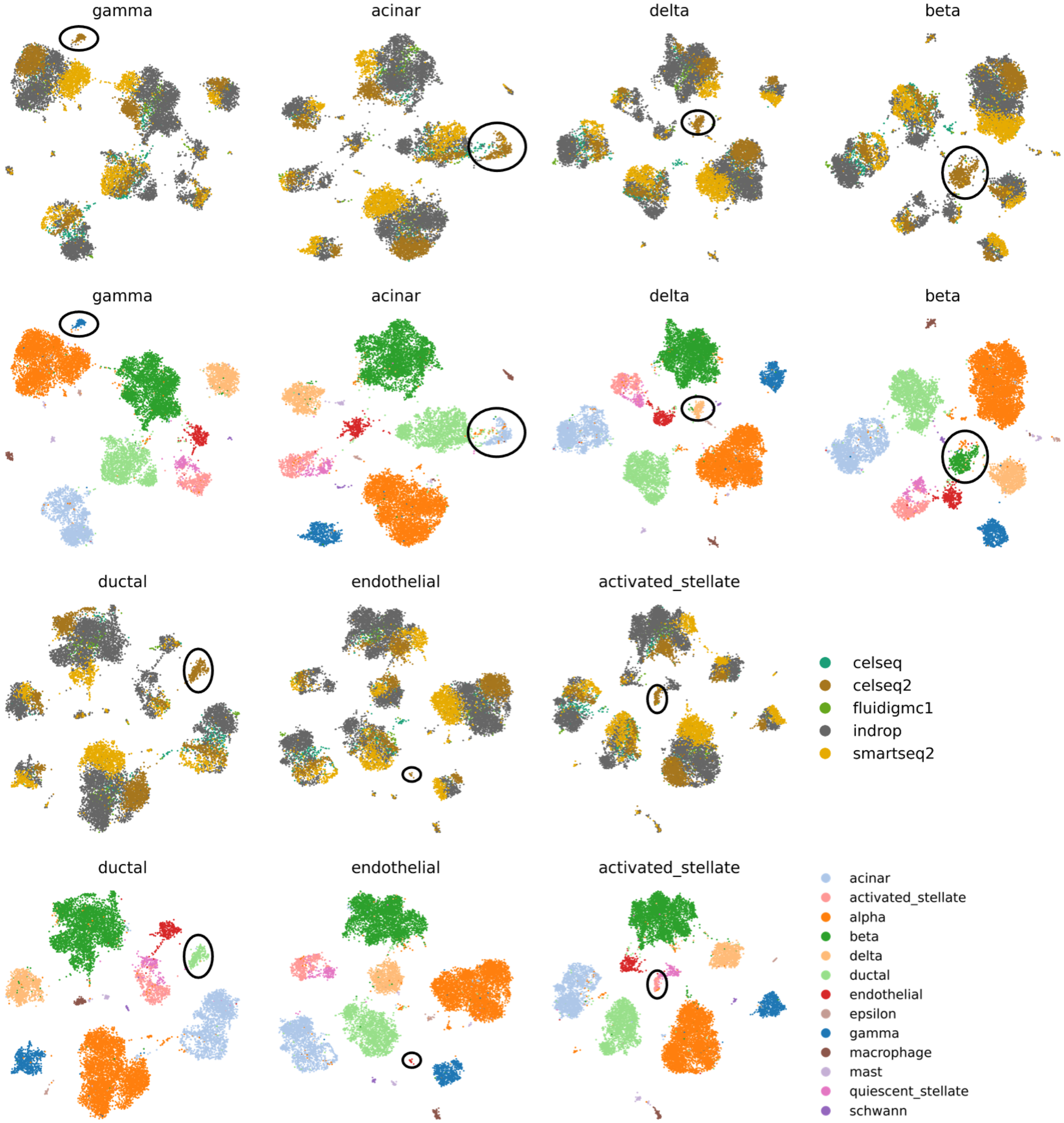


Supplementary Figure 2: Querying the pancreas islet cell reference embeddings for additional choices of held out cell types. For each plot, the corresponding held out cell type is denoted by the plot's title. For all plots, `celseq`, `fluidigmc1`, and `indrop` were used to train an HD-AE model and create a set of reference embeddings, while `celseq2` and `smartseq2` were used as query batches. Held-out cell types (black circles) were removed from all batches except `celseq2` before training the models. Plots are provided for all cell types with at least 10 cells in the `celseq2` batch.

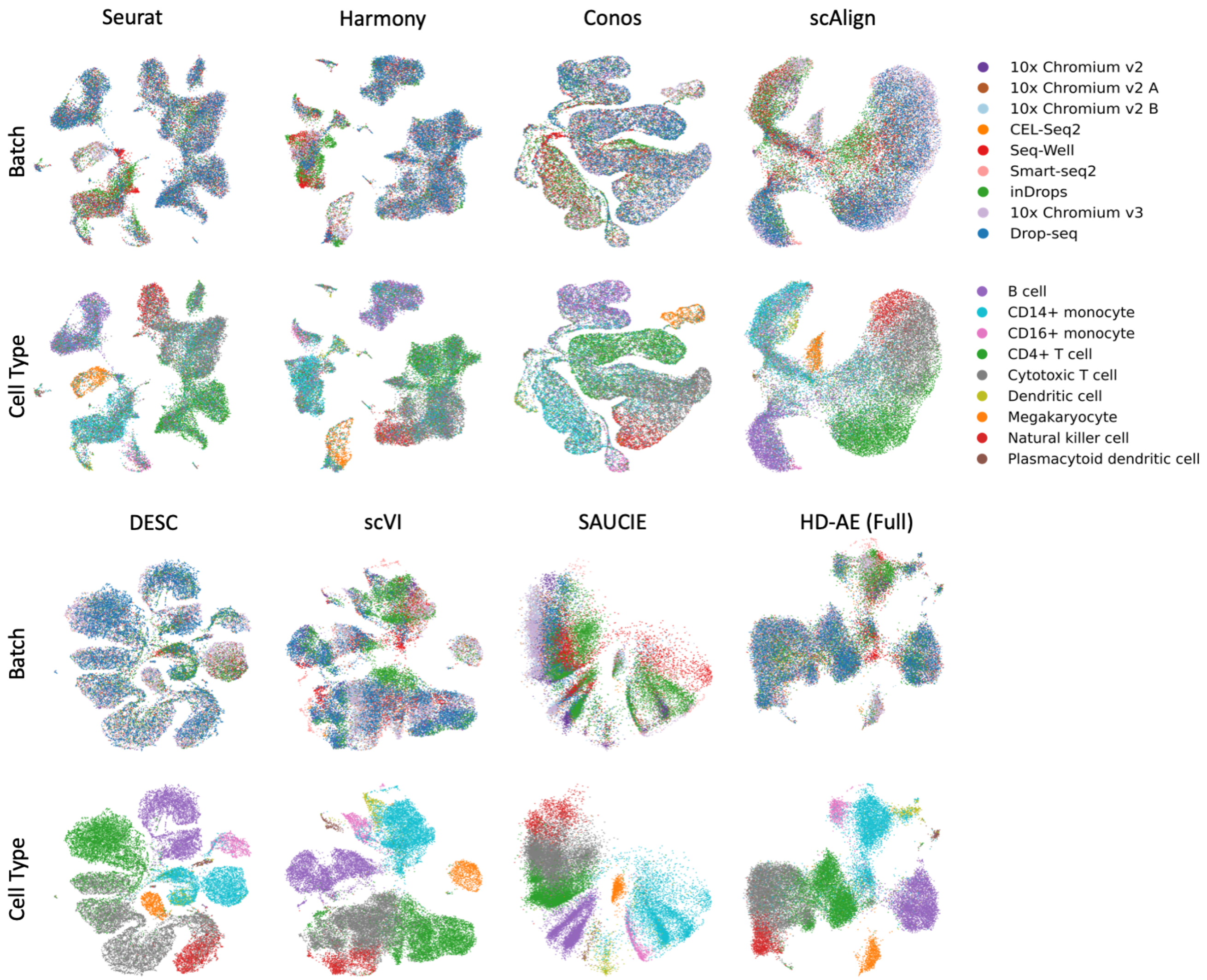


Supplementary Figure 3: Plots of PBMC embeddings for each baseline method. Because SAUCIE has a two-dimensional latent space, we plotted its embeddings directly. For all other methods, we used UMAP to generate the visualizations.

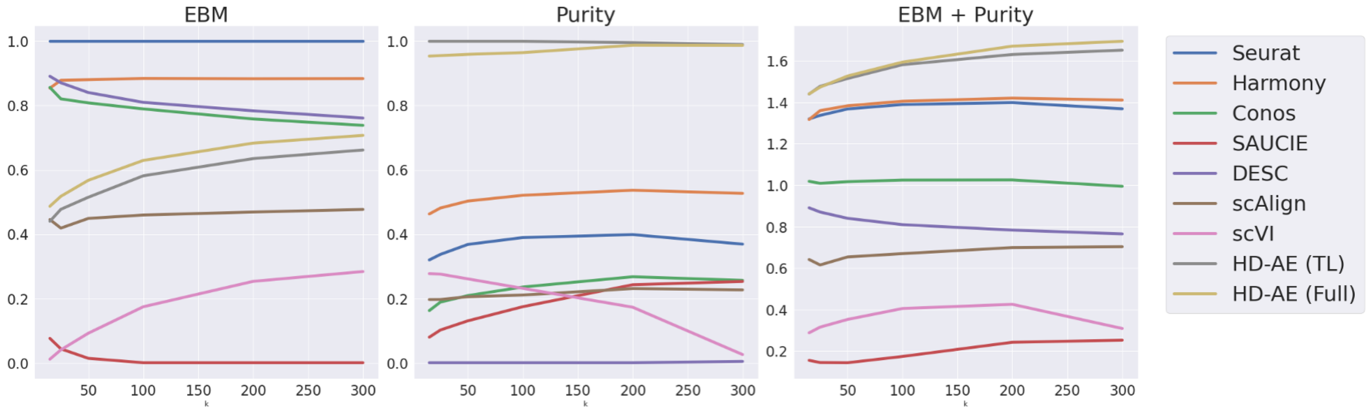


Supplementary Figure 4: Normalized entropy of batch mixing (EBM) and $k$-nearest neighbor purity scores (and their sums) for varying values of neighborhood size $k$ for HD-AE and each baseline method on the PBMC dataset.

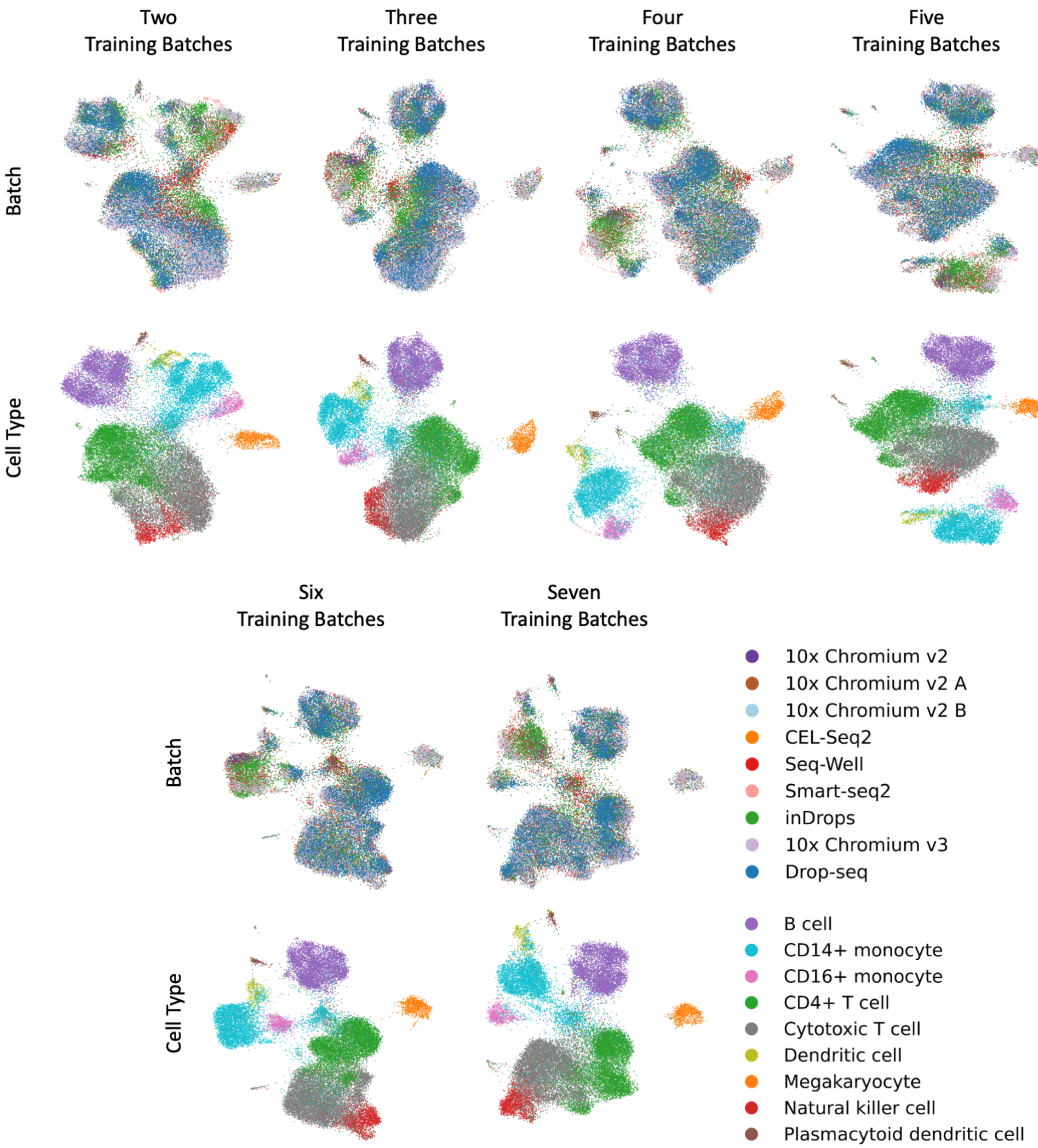


Supplementary Figure 5: UMAP plots of HD-AE embeddings of PBMC data for models trained with varying numbers of training batches.

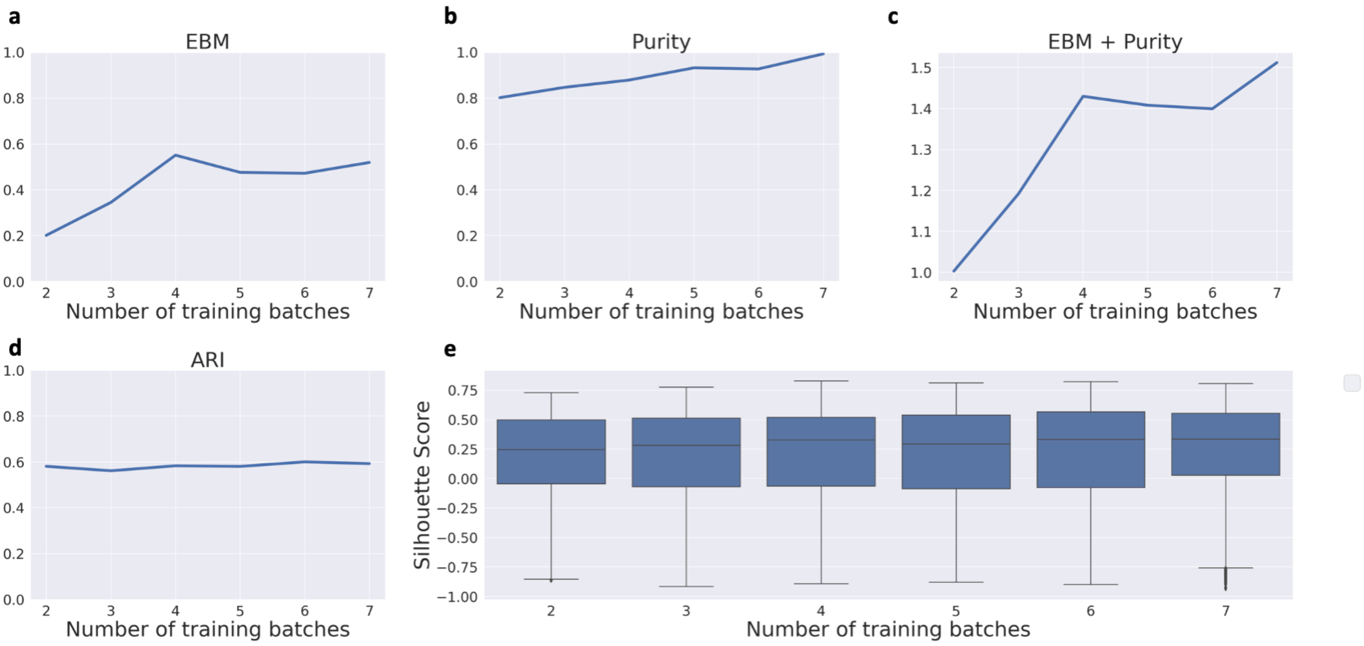


Supplementary Figure 6: **(a-c)** Entropy of batch mixing (EBM) and $k$-nearest neighbor purity for HD-AE on the PBMC dataset with varying numbers of training batches. EBM and kNN purity were normalized to lie on the same scale as in Figure **??**. **(d-e)** Adjusted Rand index (ARI) and distributions of silhouette scores for HD-AE models trained with different numbers of training batches on the PBMC dataset.

**Supplementary Note 1**: Adapting scVI, DESC, and SAUCIE for transfer learning

While scVI, DESC, and SAUCIE were not originally designed for the transfer learning setting, we could still use them for transfer learning due to their shared reliance on autoencoder-based architectures.

SAUCIE's API already supported using pretrained models to embed data from new batches at test time, so using SAUCIE for transfer learning simply involved another call to the library's `get_embedding` function. Similarly, DESC's API already supported embedding points from new batches at test time via the library's `extract_features` function.

Adapting scVI for transfer learning involved more substantial changes. scVI uses a conditional variational autoencoder to correct for batch effects. In particular, in addition to gene expression levels, batch labels are fed to scVI's encoder as an input feature. This procedure prevents a pretrained scVI model from being used for new batches at test time since it will not know how to handle samples with previously unseen batch labels. Indeed, the official scVI API throws an error when a model is fed samples from a new batch at test time. Thus, we were forced to disable scVI's batch effect correction capabilities (thereby removing the need to input batch labels to scVI's encoder) in order to use scVI for transfer learning.

**Supplementary Note 2**: Further details on DESC's and SAUCIE's loss functions and architectures

DESC is an autoencoder model trained to integrate scRNA-seq data by iteratively optimizing a clustering objective function. After pretraining the autoencoder model, the encoder network's weights are further optimized using an iterative clustering process. First, for each cell $i$ and cluster $j$, soft clustering assignments $q_{ij}$ are computed between the cell's embedding $z_i$ and the cluster's centroid $\mu_j$ using Student's t-distribution as a kernel:

$$q_{ij} = \frac{(1 + ||z_i - \mu_j||^2/\alpha)^{-1}}{\sum_{j'}(1 + ||z_i - \mu_{j'}||^2/\alpha)^{-1}}.$$

Next, these clusters are refined by minimizing the KL divergence between the soft clustering assignments $q_i$ and an auxiliary distribution $p_i$ for each cell

$$L = KL(P||Q) = \sum_{i=1}^{n} \sum_{j=1}^{K} p_{ij} \log \frac{p_{ij}}{q_{ij}},$$

where the auxiliary distribution value for each cell $i$ and cluster $j$ is defined as

$$p_{ij} = \frac{q_{ij}^2 / \sum_{i=1}^{n} q_{ij}}{\sum_{j=1}^{K} (q_{ij}^2 / \sum_{i=1}^{n} q_{ij})}.$$

Minimizing such a loss encourages the encoder network to group embeddings in tighter clusters, and we hypothesize that this behavior is detrimental for DESC's performance in the transfer learning setting. In particular, we found that embeddings of new previously unseen cell types were consistently adjacent to or mixed with clusters of cell types seen during training rather than forming separate clusters.

SAUCIE consists of an autoencoder with a two-dimensional bottleneck layer. Moreover, batch effects are corrected in SAUCIE's embedding space by penalizing the Maximum Mean Discrepancy statistic [1]. While such choices do result in mixing between batches, the resulting lack of expressive power with such a low-dimensional latent space destroys much of the structure of the original data in the embedding space; many cell types do not form contiguous clusters in the embedding space, even cells that come from the same batch of origin.

**Supplementary Note 3**: Baseline models and hyperparameter choices

For each baseline method, we used the publicly available implementations provided by the respective authors. For Seurat [2], Harmony [3], Conos [4], and scAlign [5], these implementations are available in the Batchelor (`https://bioconductor.org/packages/release/bioc/html/batchelor.html`), Seurat (`https://github.com/satijalab/seurat`), Harmony (`https://github.com/immunogenomics/harmony`), Conos (`https://github.com/kharchenkolab/`

conos), and scAlign (https://github.com/quon-titative-biology/scAlign) R packages, respectively. scVI [6], DESC [7], and SAUCIE [8] are implemented in the scVI-tools (https://github.com/YosefLab/scvi-tools), DESC (https://github.com/eleozzr/desc), and SAUCIE (https://github.com/KrishnaswamyLab/SAUCIE) Python packages, respectively.

For our HD-AD models, we used encoder networks that contained two hidden layers of size 500 and 250 with a latent space of size 50. The decoder networks followed the same structure in reverse. We optimized the models using ADAM [9] with a learning rate of 1e-3 and used 100 epochs for training. We used a minibatch size of 128 for training. We set $\lambda = 1$ for all experiments.

For Seurat, Harmony, and Conos, we used each package's default hyperparameters. To obtain embeddings for Seurat, we applied PCA to its results with 50 components. We also set the number of components to 50 for the PCA steps in Harmony and Conos.

For scVI, we used the default hyperparameters and chose the number of training epochs via early stopping with a patience parameter of 10. We initially set the latent space size to 50 (to match that of HD-AE) but found that doing so worsened performance compared to the default latent space size.

For DESC, we set the `louvain_resolution` parameter to 0.2 for our pancreas experiments and 0.6 for our PBMC experiments. These values were chosen by searching over the range $[0.1, 1]$ for both datasets and selecting the value with the best performance. As with scVI, we found that using the default latent space dimension performed better than setting it to 50. All other DESC hyperparameters kept their default values.

For scAlign, we used the default hyperparameters and chose the number of training epochs using early stopping with a patience of 10.

For SAUCIE, we set the number of epochs to 1000, and all other hyperparameters kept their default values.

**Supplementary Note 4**: Further details on the Hilbert-Schmidt Independence Criterion (HSIC)

Suppose we have two random variables $X$ and $Y$ defined over the compact domains $\mathcal{X}$ and $\mathcal{Y}$ with joint distribution $p_{XY}$. Moreover, suppose we have two separable reproducing kernel Hilbert spaces (RKHSs) $\mathcal{F}$ and $\mathcal{G}$. We denote the kernels associated with $\mathcal{F}$ and $\mathcal{G}$ as $k$ and $l$, respectively, with corresponding feature maps $\gamma$ and $\eta$. Let $||\cdot||_{HS}$ denote the Hilbert-Schmidt norm and $\otimes$ denote the tensor product. We also define $\mu_x \in \mathcal{F}$ and $\mu_y \in \mathcal{G}$ such that $\langle \mu_x, s\rangle_{\mathcal{F}} = \mathbb{E}[\langle \gamma(x), s\rangle] = \mathbb{E}[s(x)]$ and $\langle \mu_y, t\rangle_{\mathcal{G}} = \mathbb{E}[\langle \eta(y), t\rangle] = \mathbb{E}[t(y)]$; such elements are guaranteed to exist by the Riesz Representation Theorem if $k$ and $l$ are bounded. The HSIC is then defined as

$$\mathrm{HSIC}(X, Y; \mathcal{F}, \mathcal{G}) = \left|\left|\mathbb{E}_{x,y}[(\gamma(x) - \mu_x) \otimes (\eta(y) - \mu_y)]\right|\right|_{HS}^2.$$

$\mathrm{HSIC}(X, Y; \mathcal{F}, \mathcal{G}) = 0 \iff X \perp\!\!\!\perp Y$ as long as the kernels corresponding to $\mathcal{F}$ and $\mathcal{G}$ are universal kernels, a class that includes the widely used Gaussian and Laplacian kernels [10]. Moreover, if $\{(x_i, y_i)\}_{i=1}^n$ are i.i.d. samples from $p_{XY}$, the HSIC can be empirically estimated via

$$\widehat{\mathrm{HSIC}}(\{(x_i, y_i)\}_{i=1}^n) = \frac{1}{(n-1)^2}\mathrm{Tr}(KHLH),$$

where $K_{ij} = k(x_i, x_j)$ and $L_{ij} = l(y_i, y_j)$ are Gram matrices for $k$ and $l$, respectively, $H$ is a centering matrix, and Tr denotes the trace operator. $\widehat{\mathrm{HSIC}}$ converges to the true HSIC at a rate of $\mathcal{O}(n^{-1/2})$ and has bias on the order of $\mathcal{O}(n^{-1})$.
